# Habitat fragmentation selects for low dispersal in an ant species

**DOI:** 10.1101/2023.05.02.539042

**Authors:** Basile Finand, Nicolas Loeuille, Céline Bocquet, Pierre Fédérici, Joséphine Ledamoisel, Thibaud Monnin

## Abstract

Increased habitat fragmentation is one of the major global changes affecting biodiversity. It is characterised by a decrease in habitat availability and by an increase in the isolation of suitable habitat patches. The dispersal capacities of species may evolve in response to increased habitat fragmentation. Spatial heterogeneities and/or costs of dispersal, which are directly linked to habitat fragmentation, tend to select for lower dispersal abilities. We studied the effects of habitat fragmentation on dispersal using an ant species that exhibits a marked dispersal polymorphism. *Myrmecina graminicola* produces winged queens dispersing by flight over long distances, or apterous queens dispersing on foot over short distances. We sampled queens in 24 forests around Paris and 25 parks within Paris, representing varied levels of habitat fragmentation and habitat size. We identified the queen morphotypes in each environment and used it as a proxy of dispersal. Winged queens predominated in both environments. However, apterous queens were comparatively more common in parks than in forests, suggesting that high fragmentation counterselects dispersal in this species. We argue that this is because dispersing within urban environments is very costly and discuss the factors favouring each queen morph or resulting in their co-occurrence (maintenance of polymorphism).

## INTRODUCTION

Fragmentation is the modification of habitats into smaller and more isolated patches. It results in metapopulations of limited connectivity and has direct effects on species evolution and survival, by reducing gene flows and by decreasing population size [1]. Dispersal is defined as the movement of individuals associated with gene flows across space [2]. It is an important species’ trait. It can evolve, among other things, as a response to habitat fragmentation. It allows individuals to move between patches and to colonise empty ones [3]. Dispersal can therefore favour the maintenance of metapopulations [4]. However, dispersal can be selected against in some circumstances. Fragmentation increases spatial heterogeneity hence dispersal costs, as the risk of arriving in a hostile environment is largely increased, which may select for lower dispersal. This is observed in both theoretical [5–7] and empirical studies on various groups of species [8–10].

Some ant species are highly suitable study organisms to investigate the evolution of dispersal because they show a marked dispersal polymorphism [11]. Most ants either produce winged queens or apterous queens. Winged queens disperse over long distances by flight and found new colonies solitarily (a strategy called independent colony foundation, [12]). In contrast, apterous queens disperse on foot over short distances and found new colonies with the help of related workers (dependent colony foundation or colony fission). While most species produce one queen morph only, some species produce both morphs, including within populations, and are therefore an interesting model to study dispersal in fragmented landscapes. Examples include *Myrmecina graminicola* [13], *M. nipponica* [14], *Chelaner sp*. [15] and *Leptothorax canadensis* [16]. The relative abundance of each strategy varies with environmental factors, i.e. altitude in *M. nipponica* [17], habitat patchiness and isolation in *L. canadensis* [16] and drought in *Chelaner sp*. [15].

The aim of this study is to understand how dispersal evolves depending on habitat fragmentation. Specifically, we test the hypothesis that dispersal is selected against in highly fragmented habitats. We use the ant *M. graminicola* as a model organism because it exhibits a marked, genetically determined, dispersal polymorphism between winged and apterous queens [18]. We sampled queens in variously fragmented habitats (25 urban Parisian parks and 24 forests around Paris) and uncovered their dispersal strategies. We expected to find a higher proportion of apterous queens in urban parks than in forests, due to the higher spatial heterogeneity of urban environments that leads to higher risks of dispersal into an unfavourable habitat. In addition, we expected apterous queens to be more common in small parks and forests than in large parks and forests, because of the higher habitat fragmentation.

## MATERIAL AND METHODS

### The model species, Myrmecina graminicola

*M. graminicola* (Latreille 1802) is a western palearctic ant species largely distributed in Europe [19]. It produces winged queens and apterous queens, the first type showing high dispersal capacities and independent colony foundation, while the second uses colony fission and disperses at small distances [13]. This polymorphism is genetically based [18]. This species lives in relatively warm and damp forested habitats, and nests in the soil or beneath rocks or moss [13,20]. It forages in the leaf litter and feeds on small arthropods.

### Site selection

Sampling was carried out in 25 urban parks in Paris and 24 forests around Paris (in the Île de France region). Parks are heterogeneous habitats for this species, with favourable wooded areas or groves intertwined with inhospitable areas such as open lawn and paths. In addition, each park is surrounded by a totally inhabitable environment (streets and buildings of a very dense city). This clear dichotomy between parks and their surrounding matrix allows a clear categorisation of parks according to their size (below). In contrast, forests are less heterogeneous habitats, with only a few paths interrupting ubiquitous favourable wooded areas. In addition, most forests are surrounded by less favourable yet habitable environments such as farmland and villages with many tree borders. Therefore, forests were not categorised by their size but by their level of fragmentation.

Forests (n = 24) were sampled in 2020. They were categorised into three fragmentation classes (Figure 1a, Table 1). Highly fragmented forests (n =6) had less than 5% of forest cover (measured on GIS maps) within a 1 Km radius circle around the sample point (i.e. were isolated forest patches). Within the same range, medium fragmented forests (n = 8) had between 30 and 70% of forest cover, and lowly fragmented forests (n = 10) had between 75 and 95% of forest cover (i.e. were large continuous forests). Forests were selected prior sampling using the database BD Forêt® Version 2.0 from the *Institut national de l’information géographique et forestière* of France (IGN, https://inventaire-forestier.ign.fr/carto/afficherCarto/V2, last visit: 19/07/2022). We selected forests so that forest classes were evenly distributed around Paris (Figure 1a), and forests were well separated from one another hence considered independent sampling points (the nearest forests were 6.2 Km apart).

**Figure 1:**
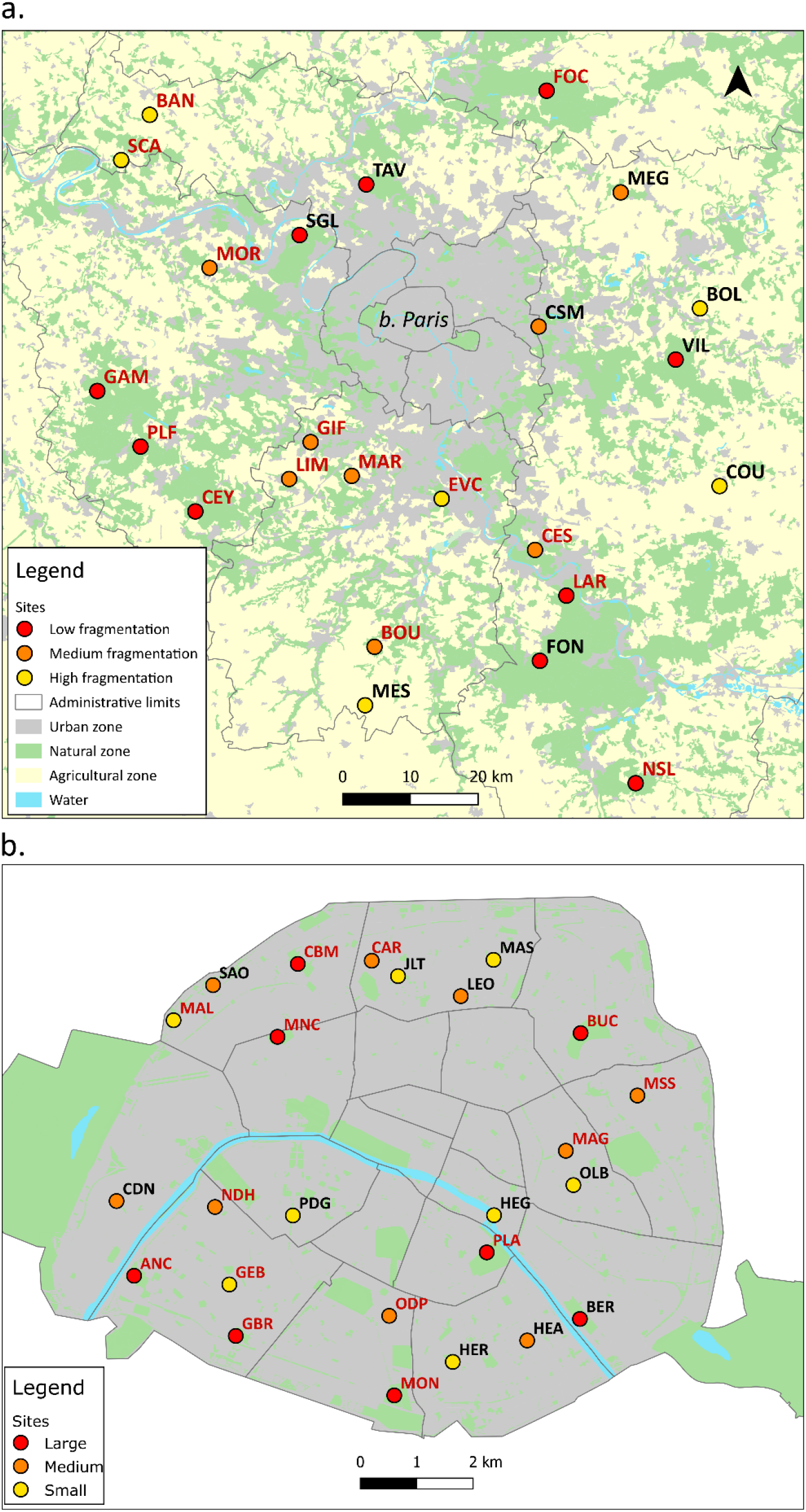
Map of the sampling sites in (a) forests around Paris (Ile-de-France region) and (b) parks in Paris. The colour of the circles indicates the fragmentation level of forests or size of parks. Red labels indicate sites where queens were found.

**Table 1:** Number of each queen morph across sites

| **Site** | **Habitat** | **Fragmentation**<br>**or Size** | **Number of**<br>**workers** | **Number of**<br>**apterous queens** | **Number of**<br>**winged queens** |
| --- | --- | --- | --- | --- | --- |
| CEY | Forest | Low | 105 | 0 | 4 |
| FOC | Forest | Low | 229 | 2 | 4 |
| FON | Forest | Low | 9 | 0 | 0 |
| GAM | Forest | Low | 278 | 0 | 8 |
| LAR | Forest | Low | 192 | 5 | 15 |
| NSL | Forest | Low | 19 | 0 | 1 |
| PLF | Forest | Low | 49 | 0 | 3 |
| SGL | Forest | Low | 0 | 0 | 0 |
| TAV | Forest | Low | 0 | 0 | 0 |
| VIL | Forest | Low | 0 | 0 | 0 |
| BOU | Forest | Medium | 69 | 0 | 1 |
| CES | Forest | Medium | 216 | 0 | 6 |
| CSM | Forest | Medium | 111 | 0 | 0 |
| GIF | Forest | Medium | 255 | 0 | 27 |
| LIM | Forest | Medium | 78 | 0 | 7 |
| MAR | Forest | Medium | 36 | 0 | 2 |
| MEG | Forest | Medium | 11 | 0 | 0 |
| MOR | Forest | Medium | 247 | 0 | 5 |
| BAN | Forest | High | 119 | 0 | 1 |
| BOL | Forest | High | 7 | 0 | 0 |
| COU | Forest | High | 3 | 0 | 0 |
| EVC | Forest | High | 56 | 0 | 1 |
| MES | Forest | High | 19 | 0 | 0 |
| SCA | Forest | High | 37 | 0 | 8 |
| ANC | Park | Large | 197 | 0 | 2 |
| BER | Park | Large | 41 | 0 | 0 |
| BUC | Park | Large | 827 | 1 | 2 |
| CBM | Park | Large | 75 | 0 | 6 |
| GBR | Park | Large | 55 | 0 | 5 |
| MNC | Park | Large | 212 | 1 | 2 |
| MON | Park | Large | 90 | 1 | 0 |
| PLA | Park | Large | 1008 | 2 | 0 |
| CAR | Park | Medium | 20 | 0 | 1 |
| CDN | Park | Medium | 79 | 0 | 0 |
| HEA | Park | Medium | 39 | 0 | 0 |
| LEO | Park | Medium | 24 | 0 | 0 |
| MAG | Park | Medium | 370 | 2 | 4 |
| MSS | Park | Medium | 38 | 0 | 1 |
| NDH | Park | Medium | 45 | 1 | 1 |
| ODP | Park | Medium | 19 | 1 | 1 |
| SAO | Park | Medium | 90 | 0 | 0 |
| GEB | Park | Small | 263 | 3 | 2 |
| HEG | Park | Small | 1 | 0 | 0 |
| HER | Park | Small | 1 | 0 | 0 |
| JLT | Park | Small | 37 | 0 | 0 |
| MAL | Park | Small | 74 | 0 | 1 |
| MAS | Park | Small | 152 | 0 | 0 |
| OLB | Park | Small | 23 | 0 | 0 |
| PDG | Park | Small | 139 | 0 | 0 |

Parks (n = 25) were sampled in 2021. They were categorised into three size classes (Figure 1b, Table 1). Small parks (n = 8) ranged from 500 to 2 000 m², medium parks (n = 9) ranged from 5 000 to 12 000 m², and large parks (n = 8) ranged from 80 000 to 250 000 m². All parks are managed by the City of Paris Council, except the botanical garden (*Jardin des plantes*) which is managed by the National Museum of Natural History. We carefully selected parks so that size classes were spread throughout the city and that parks were well separated from one another to avoid pseudoreplication.

### Sampling

Forests were sampled between May and July 2020, between 11am and 4pm, on rainless days. We laid out a 100 m long transect, along which 12 quadrats of one square meter each were positioned equally spaced (every 9 m). Considering the small colony size and worker size of *M. graminicola*, each quadrat can be considered an independent sampling point in that individuals from a given colony were unlikely to be present on two quadrats. For each forest, we collected the litter and superficial soil from each of the 12 quadrats and extracted the organisms therein contained, using the Winkler extraction method [21] (Figure S2) as detailed below.

Parks were sampled between May and August 2021, between 11am and 4pm, also on rainless days only. Due to the spatial configuration of the parks, it was neither possible nor adequate to sample along a linear 100 m long transect as in forests. Instead, we purposefully placed the 12 quadrats in the closed environments of the park, i.e. under trees or bushes providing shade and producing leaf litter. One park (OLB) was so small that only five quadrats could be placed and sampled, and owing to a disturbance only ten quadrats were sampled in another park (MAS). We sampled the litter and superficial soil in parks as we did in forests.

*M. graminicola* forages in the leaf litter and has small and slow-moving workers. Sampling was thus carried out using the Winkler extraction method [21] (Figure S2). For each quadrat, we recovered the litter and the first centimetre of soil and immediately sifted this using 1 cm² sieves. This allows removing leaves, twigs and stones while retaining the soil and its organisms including *M. graminicola*. The sifted litter from each quadrat was kelp in a large plastic bag (n = 12) until returning to the laboratory. Organisms were extracted by moving the litter to sieve bags that were hanged in a Winkler extractor (n = 12) for 48 hours. As the litter gradually dried out, organisms moved away and fell into an alcohol-filled collecting vial [21]. This method allowed sampling foragers of *M. graminicola*, as well as many queens because nests of this species are shallow. Collecting queens is necessary given that it is their thorax morphologies that allows identifying their dispersal strategies.

### Characterisation of queen morphs

The Winkler extractors yielded alcohol-filled collecting vials containing some fine litter and large numbers of organisms. We separated ants from other organisms at the naked eye and under a binocular loupe. We then identified and put aside *M. graminicola* individuals using a taxonomic guide (21). We separated queens from workers, the former having bigger bodies and eyes, and identified queen morphs (winged/apterous) following [13]. The two queen morphs have the same size but differ in thorax morphology. Winged queens have a typical ant queen thorax morphology. They remove their wings after dispersal and their thorax bears scars at the point of insertion of the wings. Apterous queens have a different thorax morphology and bear no such marks as they are born wingless (see details in [13]).

### Analyses

The analyses were done with R 3.6.2 [22]. We compared the number of queens of each morph between the two habitat types, and between each patch size class within each habitat type, with Fisher’s exact tests. Additionally, we analysed the differences between sites by comparing the proportion of each queen morph (i.e. relative abundance of each dispersal strategy) in each site as a function of habitat type and patch size with a binomial GLM. We checked the conditions of application visually and statistically (Shapiro test).

## RESULTS

*M. graminicola* was one of the most abundant ant species in our samples. We collected 140 queens in 15 forests (in six large, six medium and three small forests) and 14 parks (in seven large, five medium and two small parks) (Table 1). We decided to pool the data from small and medium parks because of the relatively low abundance of queens in them (i.e. for statistical analysis), and because of their relative similarity in size as compared to large parks (see Figure S1).

Queens were collected in as many forests than parks (15 out of 24 forests vs 14 out of 25 parks, Figure 2a). As predicted, apterous queens occurred more frequently in the (more fragmented) urban habitats. They were collected in eight of the 14 parks vs only two of the 15 forests (Fisher’s exact test, p=0.02, Figure 2a). In addition, apterous queens were present in all park sizes but only in lowly fragmented forests. Considering queen abundance rather than occurrence shows a similar pattern. Apterous queens were more abundant in the (more fragmented) urban habitats, with 30% of queens having the apterous morphology in parks versus 7% in forests (Fisher’s exact test, p=0.0008, Figure 2b). Note that this relative abundance of queen morphs in forests is corroborated by an active colony search carried out in addition to Winkler extraction, that yielded an additional 94 queens with a similar morph distribution (Figure S2). Apterous queens were equally abundant in parks of all sizes (22.7% and 38.9% of queens are apterous in large and medium+small parks, respectively, Fisher’s exact test, p=0.32), whereas they were only found in large forests (Fisher’s exact test, p=0.0056) where they represented 16.7% of queens (Figure 2b).

**Figure 2:**
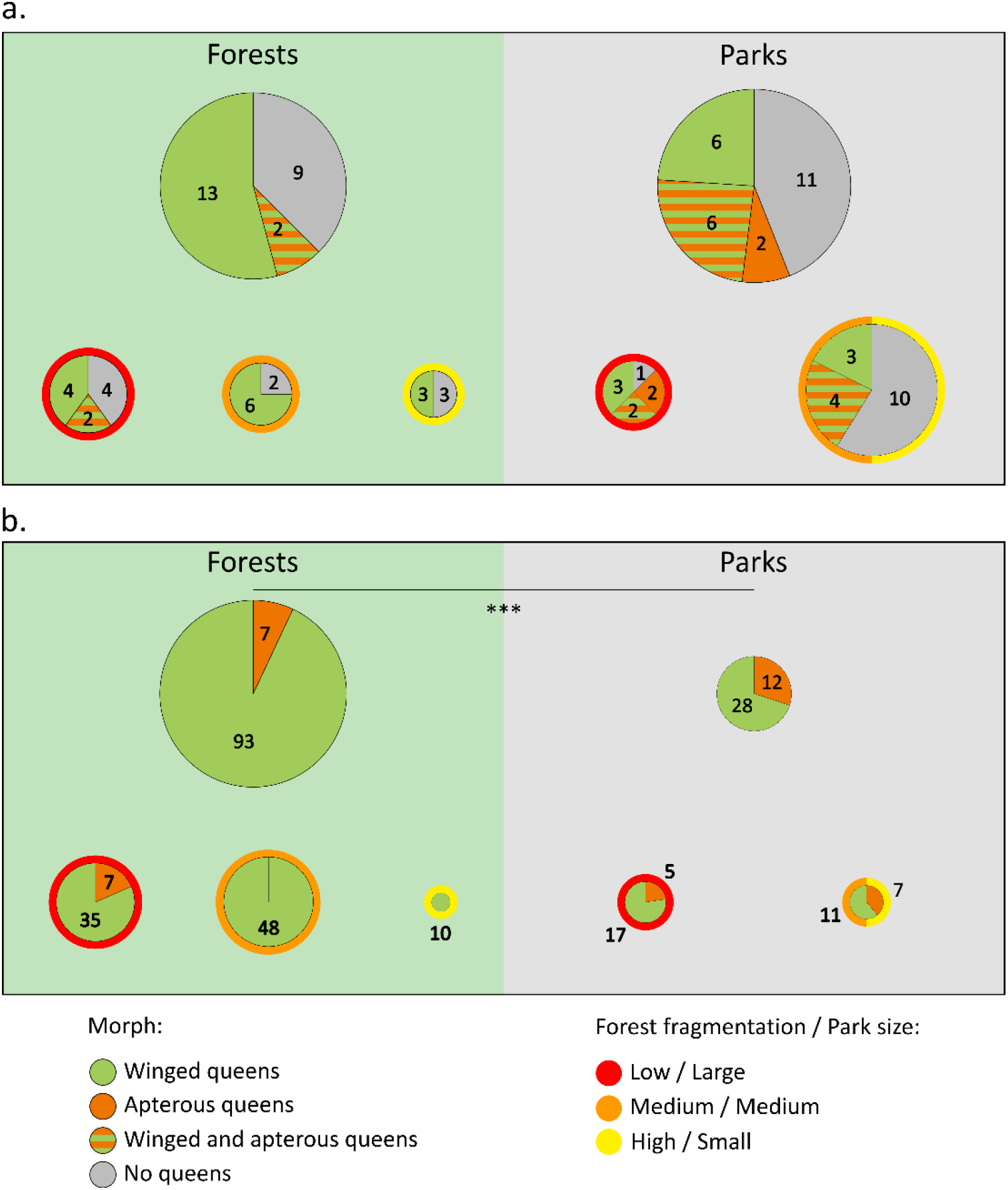
Occurrence (a) and abundance (b) of queen morphs in forests and parks. Pie chart size is proportional to sample size. Note that small and medium parks were pooled (see Fig. S1). Panel (a) shows the number of sites where each morph (winged vs apterous) was found, i.e. their occurrence, per habitat type (forests vs parks, top) and decomposed according to forest fragmentation level or park size (bottom). Apterous queens occurred in two large forests only while they occurred in eight parks. They typically co-occurred with winged queens. Panel (b) shows the number of queens of each morph i.e. their abundance, per habitat (top) and fragmentation level or size (bottom). Apterous queens were more abundant in parks than in forests (Fisher’s exact test, p=0.0008).

The analysis above compares site types (forests vs parks) but does not show the variability of morphotype distributions, which is shown in Figure 3. Apterous queens represent 32.9 +/- 36.0% of queens in parks versus 3.9 +/- 10.4% in forests (GLM, deviance = 11.59, p < 0.001, figure 3a). Among forests, apterous queens were only found in large forests (GLM, deviance = 12.88, p = 0.002), with a proportion of 9.7 +/- 15.3% of the queens (Figure 3b). On the contrary, apterous queens were equally abundant in parks regardless of their size (38.1 +/- 44.8% and 27.6 +/- 27.0% of queens are apterous in large and medium+small parks, respectively, GLM, deviance = 1.23, p = 0.267, Figure 3c).

**Figure 3:**
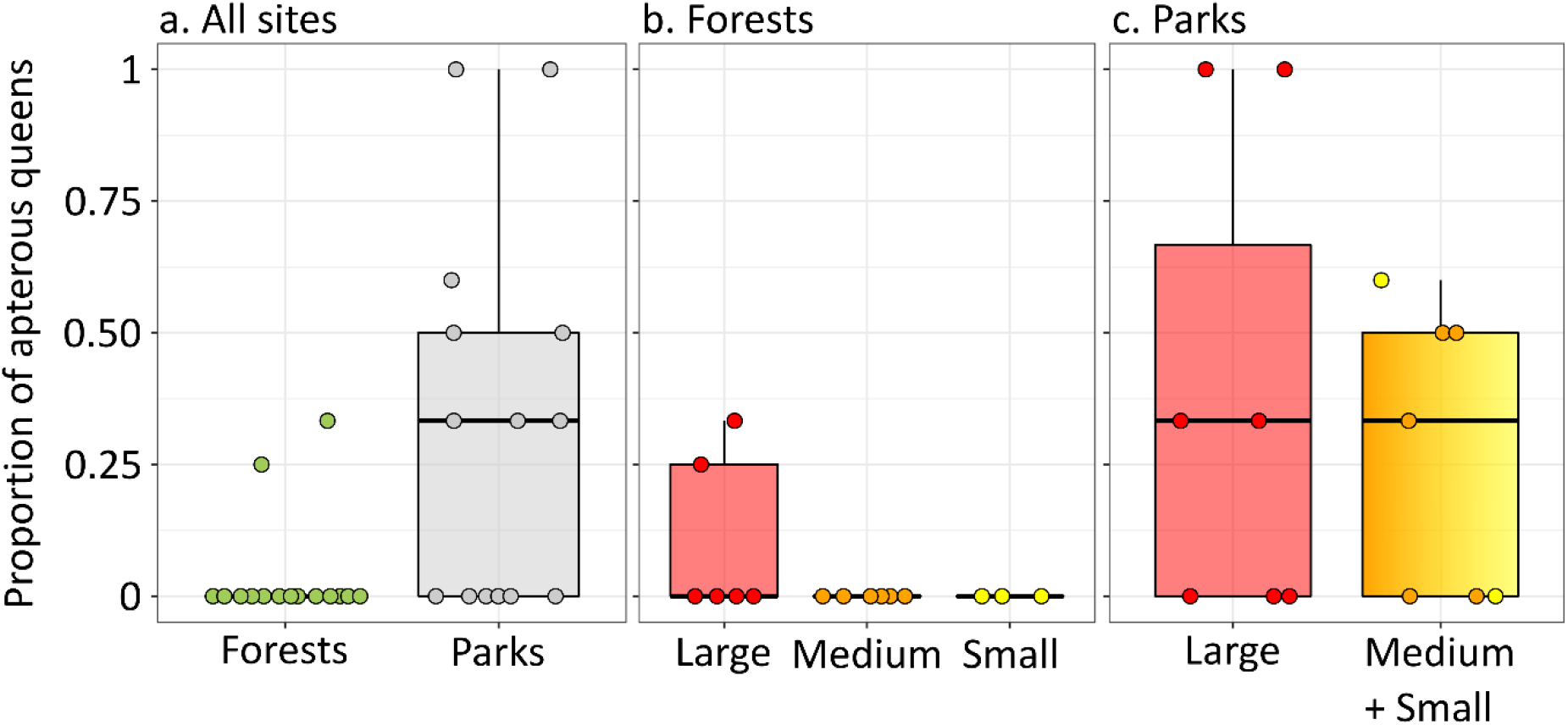
Proportion of apterous queens according to (a) habitat types, and (b, c) habitat type and fragmentation/size. Note that small and medium parks were pooled (see Fig. S1). Apterous queens were more abundant in parks than in forests (GLM, deviance = 11.59, p < 0.001), and among forests they were more abundant in large forests than in medium or small forests (GLM, deviance = 12.88, p = 0.002).

## DISCUSSION

Multiple forces influence the evolution of dispersal, and the study of species with dispersal polymorphism can provide a better understanding of this evolution. Our study suggests an evolutionary response of an ant species with a marked genetic-based dispersal polymorphism to habitat fragmentation. We observe more apterous queens in parks (high fragmentation) than in forests (low fragmentation). Park size had no effect on apterous queens’ abundance. In contrast, while forests with high and medium-levels of fragmentation lacked apterous queens, low fragmented forests did harbour apterous queens, albeit at a lower level than parks. This pattern may be explained by the predominance of alternative evolutionary forces in the various environments, with parks selecting against dispersal and, possibly, large forests selecting for competitiveness.

Parks are fragmented habitats. They are heterogeneous, with some habitats favourable to *M. graminicola* (closed and humid environments) intertwined with less favourable habitats (open lawn, closed bush, flower patches, paths, etc), and they are surrounded by highly inhospitable city environments (streets, buildings) where survival of newly dispersed queens and the foundation of new colonies is unlikely. This likely selects against long distance dispersal, especially given that ant queens are relatively poor fliers. Because dispersal polymorphism is genetically mediated in *M. graminicola* [18], our study provides a new animal case of fragmented habitat adaptation through dispersal evolution. This counterselection of dispersal in fragmented habitat supports our hypothesis that fragmentation, by creating spatial heterogeneities and increasing the costs of dispersal, disfavours more dispersive strategies [5– 7]. This effect of dispersal costs associated with spatial heterogeneities is congruent with theoretical works [5–7,23] as well as with empirical observations in other systems. In another ant species, *L. canadensis*, apterous queens were more abundant in small and isolated forests than in larger forests [16]. Similar results were obtained for the bug fritillary butterfly *Proclossiana eunomia* [8], the dune wolf spider *Pardosa monticola* [9] or the weed *Crepis sancta* [10], with fewer dispersal events in more fragmented habitats for the former, fewer dispersal behaviour of tiptoe for the second and a higher proportion of heavy non-dispersing seeds for the latter.

In contrast to parks, large forests essentially consist of habitats that are favourable to *M. graminicola*. Consequently, there is little risk to disperse to unfavourable environments and this selects for dispersive strategies in forests. However, we found a few apterous queens, but in large forest patches only. A possibility, assuming a trade-off between competitive ability and dispersal, would be that large favourable habitats select for competitive strategies, and therefore for low dispersal as a by-product [24,25]. Indeed, a laboratory experiment shows that *M. graminicola* colonies founded by a queen plus two or four workers survive more and grow faster than colonies founded by a solitary queen (pers com). This is congruent with several studies that show that in case of polymorphism, the competitive strategy occurs in large patches [23,26,27]. However, seven apterous queens only were found, and in two forests only, hence caution is required. Indeed, an alternative explanation is that queen polymorphism is maintained in all populations, with a low abundance of apterous queens in all forests, and that we found some in two large forests by chance. Queen morphs are genetically mediated in *M. graminicola* [18] and the development of genetic markers would allow us to dramatically increasing sample size by using the many workers sampled instead of the relatively few queens.

High fragmentation usually selects against dispersal due to increased spatial heterogeneities, as supported by our results, yet other forces can have the opposite effect of a selection for higher dispersal with increased fragmentation. For instance, fragmentation may increase inbreeding and kin competition due, in part, to the reduction in habitat size. In addition, if fragmentation varies over time this temporal variation of the environment can select for higher dispersal as a bet-hedging strategy. These forces select for more dispersal as shown in several theoretical [7,26,28–32] and empirical works [33,34].

Our study reveals that habitat fragmentation selects for queen morphology in the ant *M. graminicola*. In particular, parks harbour more apterous queens than forests, even though winged queens remain predominant in both habitats. Polymorphism is therefore maintained both in urban and forest environments. The limited dispersal of apterous queens could threaten the maintenance of the metapopulation by reducing genes flows and increasing inbreeding, thereby making each patch more sensitive to extinction [4]. However, both apterous and winged queens produce winged males, and queens can mate with males produced by either queen morphs [18], which allows some gene flow. Given the spatial variation of morphotypes we observe, this gene flow is however not sufficient to compensate for local adaptation in urban environments. The limited dispersal of apterous queens also prevents the recolonization of empty patches where the population has gone extinct [3]. Last, it decreases the capacity of species to follow the conditions that match their ecological niches, which may be particularly detrimental given the current global changes. Other factors may be at play, such as passive dispersal of *M. graminicola* to and between parks by gardeners [35]. Given the polymorphism of dispersal in this species, *M. graminicola* could be a particularly suitable system to study the evolution of dispersal in a changing environment and its consequence for metapopulation maintenance.

## Supporting information

Supplementaty informations

## ACKNOWLEDGMENTS

We thank the city of Paris and the National Museum of Natural History for allowing us to sample in Parisian parks.

## FUNDING

BF was funded by the French ministry of higher education, research and innovation.

## AUTHOR CONTRIBUTIONS

BF, TM and NL conceived the ideas and designed methodology; BF, CB, PF, TM, NL and JL collected the data; BF analysed the data; BF, NL and TM wrote the manuscript. All authors gave final approval for publication.

## DATA ACCESSIBILITY

Data and codes will be available on zenodo after revisions.

## Notes

### Competing Interest Statement

The authors have declared no competing interest.

