## Supplementaty informations for "Habitat fragmentation selects for low dispersal in an ant species"

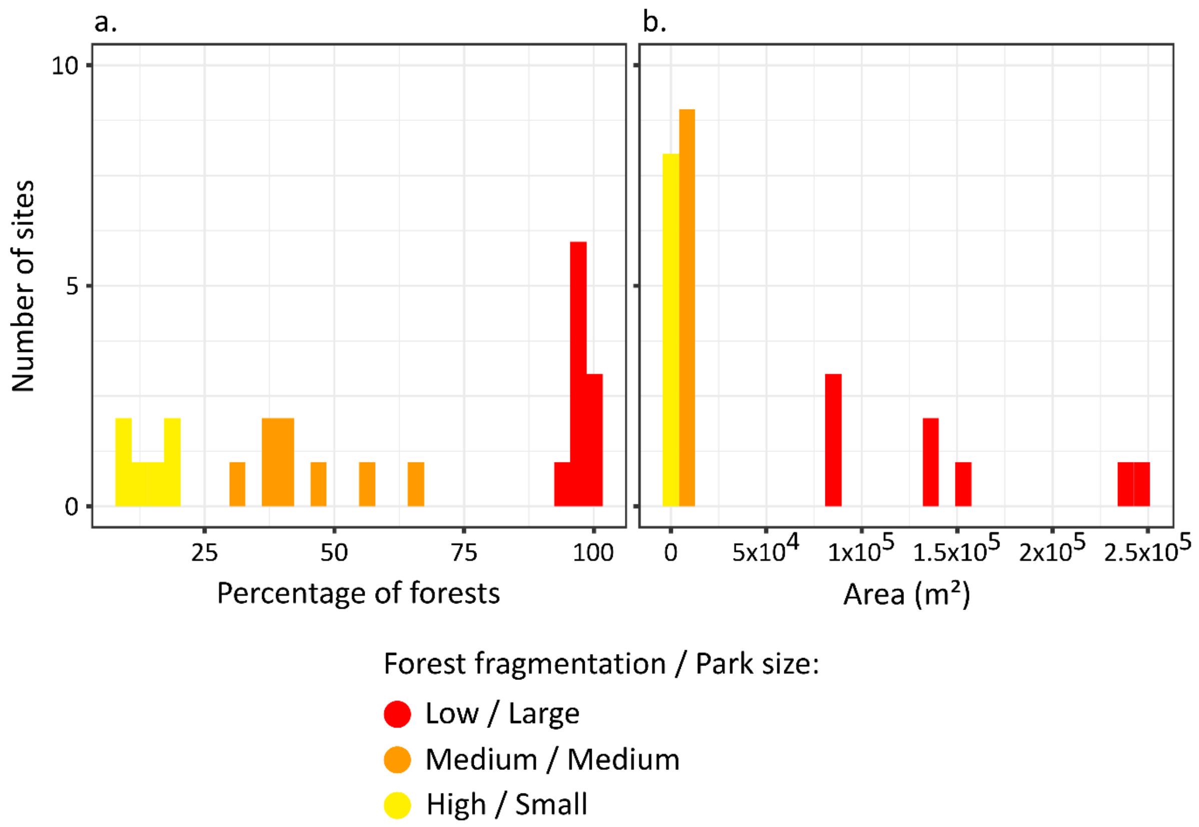


**Figure S1:** Variability of (a) fragmentation levels among the three forests classes, and (b) park size among the three park classes. Small and medium parks were pooled for analysis because (1) they yielded few queens only and (2) although medium parks (5 000 to 12 000 m²) were more than twice as large as small parks (500 to 2 000 m²) these two categories were similar in size when compared to large parks (80 000 to 250 000 m²).

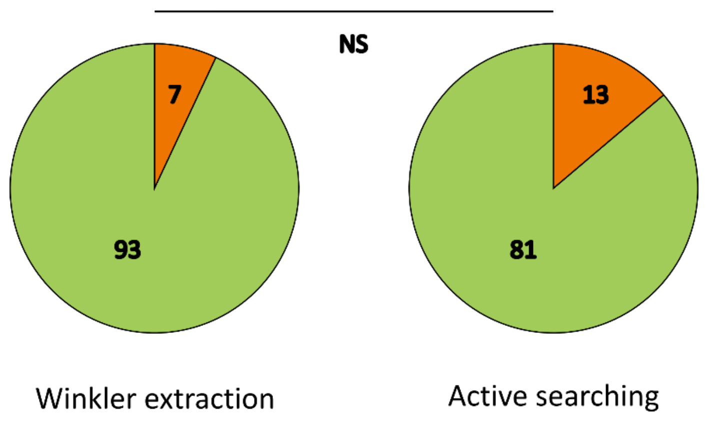


**Figure S2:** The number of winged (green) and apterous (brown) queens collected in forests by Winkler extraction and by active search did not differ (Chi-square = 1.762, p = 0.184). Active search for nests consisted in digging for a few cm under mosses, stones and half-buried broken branches, or even randomly in the soil. It was typically carried out by two persons for one hour, or sometimes by three persons for 40 min. Active search was carried out in all forests but not in parks because it was unsuccessful in the first parks sampled.

a.

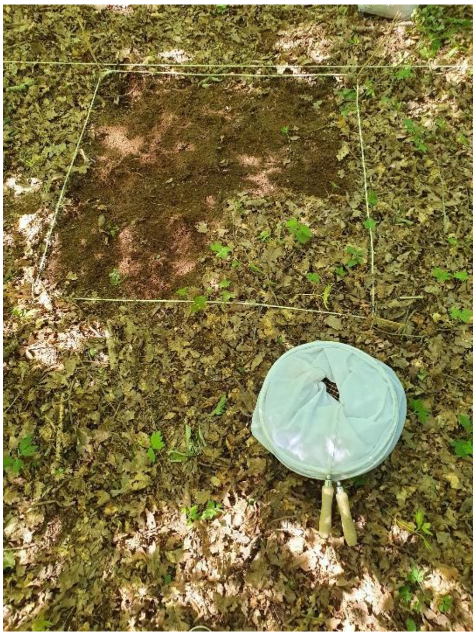

b.

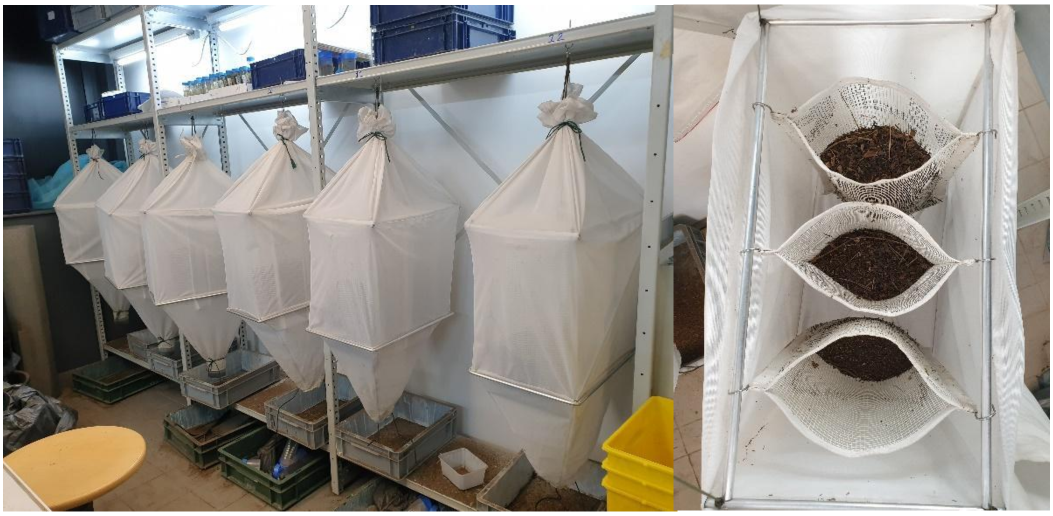

**Figure S3:** Pictures of (a) a quadrat being sifted and (b) the Winkler devices in the laboratory.

Credits: Basile Finand
